# Dog shelter facilitated transmission and maintenance of benzimidazole resistance-associated alleles in canine hookworms

**DOI:** 10.64898/2026.11.11.765975

**Authors:** Analise McDonald, Shaun Yii Woo, Michael P. Ward, Nicolle Kirkwood, Jan Šlapeta

## Abstract

Anthelmintic resistance in canine hookworms is increasingly recognised; however, environmental settings that maintain and disseminate resistance-associated alleles remain poorly defined. We hypothesised that urban dog shelters act as hotspots for transmission of benzimidazole resistance-associated alleles in *Ancylostoma caninum*. Residual faecal samples (n=173) were collected from a Sydney, Australia metropolitan shelter over four sampling periods and analysed using coprodiagnostic methods and deep amplicon sequencing targeting ITS-2 and isotype-1 β-tubulin loci. Hookworm eggs were detected in 41% of samples, with *A. caninum* identified in 97% of sequenced infections and *U. stenocephala* in 8%. The isotype-1 β-tubulin F167Y mutation was detected in 83% of *A. caninum*-positive samples, although high allele frequencies (>70%) were uncommon. Longitudinal sampling identified ten dogs that acquired *A. caninum* during shelter residence, nine of which acquired populations carrying the F167Y mutation. These findings support the role of dog shelters as hotspots for maintenance and dissemination of benzimidazole resistance-associated alleles in canine hookworms. While the impact of this dissemination remains uncertain, ongoing surveillance and targeted anthelmintic stewardship are likely to be important in high-density dog populations.

## 1. Introduction

Anthelmintic resistance in companion animal hookworms was historically considered unlikely due to the presumed maintenance of refugia, supporting routine prophylactic deworming even in intensively managed groups. However, recent evidence from North America has confirmed resistance in canine hookworm (*Ancylostoma caninum*) to benzimidazoles (BZs), tetrahydropyrimidines (pyrantel) and macrocyclic lactones (MLs), and from Australia to BZs and pyrantel (Abdullah et al., 2025; Dale et al., 2024; Geary et al., 2025; Jimenez Castro et al., 2019; Kitchen et al., 2019; Venkatesan et al., 2023). BZ-resistance is a result of well characterised genetic mutations in the isotype-1 β-tubulin of *A. caninum* that enables screening for BZ-resistance across infected dog populations (Abdullah et al., 2025; Jimenez Castro et al., 2025; Leutenegger et al., 2024; Venkatesan et al., 2023). For MLs and pyrantel resistance, the genetic targets remain elusive.

Elevated frequency of F167Y mutation in the isotype-1 β-tubulin of *A. caninum* correlates with increased BZ (thiabendazole) IC₅₀ values in egg hatch assays, supporting a genotype-phenotype relationship (Abdullah et al., 2025). However, the epidemiological context that maintain and disseminate hookworms carrying BZ-resistant alleles remain speculated or inferred (Dini et al., 2026; Geary et al., 2025; Kaplan et al., 2026; Stocker et al., 2026; Stocker and Šlapeta, 2025). Households typically containing one or a few dogs and activities such as walking dogs in the park or use of off-leash areas are unlikely to represent major drivers because hookworm infections are relatively infrequent in these populations. Instead, we hypothesised that high-density, multi-dog environments with continual introduction and turnover of animals act as transmission ‘hotspots’ in which BZ-resistant hookworms are maintained and spread among dogs.

Urban high density shelter environments are characterised by high faecal contamination, moisture-retaining substrates, crowding, high turnover and unknown treatment histories (Dale et al., 2024; Raza et al., 2018). This has been thought to assist the transmission cycle of *A. caninum* and even emergence of resistant populations (Dini et al., 2026). Furthermore, dogs exiting shelters exposed to and subsequently carrying *A. caninum* resistant alleles facilitate mixing of parasite populations (Abdullah et al., 2025; Dale et al., 2024). Therefore, shelters are plausible urban ‘hotspots’ and convergence points for the maintenance and transmission of *A. caninum* for dogs rehomed within and outside metropolitan areas (Abdullah et al., 2025; Dale et al., 2024; Raza et al., 2018).

The aim of this study was to determine whether metropolitan dog shelters act as convergence points for the maintenance and transmission of BZ-resistance associated alleles in *A. caninum* populations destined for rehoming. We have undertaken a longitudinal study within a metropolitan shelter and applied deep amplicon sequencing targeting ITS-2 and isotype 1 β-tubulin genes. We determined hookworm species composition (*A. caninum* and *U. stenocephala*) in faecal samples and frequency of detection of BZ-resistance associated SNPs (F167Y and Q134H) across and within sampled populations. Putative clinical drug failure was considered if BZ resistant allele (F167Y) exceeded 70% within a single sample. Acquisition of hookworms with resistant alleles within a shelter was demonstrated by a change of hookworm status in dogs that were sampled over multiple sampling periods.

## 2. Materials and Methods

### 2.1. Sample collection

The use of residual diagnostic canine faecal samples was in accordance with the University of Sydney (Australia) Animal Ethics Committee Protocol 2026/AE000002. Faecal samples were obtained from registered veterinary practitioners under the provision of the Veterinary Practice and Animal Welfare Acts and submitted for routine diagnostic testing to inform clinical decision-making. All samples were processed at the Veterinary Pathology Diagnostic Services (VPDS, The University of Sydney, New South Wales, Australia). The faecal samples originated from a single animal shelter facility in Western Sydney. Dogs were housed across three separate buildings within the facility (Shelters 1-3) and shared common outdoor exercise runs. All material used for hookworm species identification and assessment of BZ genetic susceptibility was de-identified prior to analysis. Breed and age were retained, but dog and shelter identities were replaced by arbitrary identifiers.

### 2.2. Coproanalysis and faecal egg counts for hookworm recovery

Faecal egg counts (FECs) were conducted using saturated salt (NaCl, specific gravity = 1.21) solution and a Whitlock Universal McMaster slide with four counting chambers and recorded as egg per gram. Egg concentration and collection from 3g of faeces was performed via centrifugal flotation (800×g for 5 min.) in 50 mL Falcon tubes containing saturated NaCl, as previously described (Abdullah et al., 2025). Briefly, the upper 10 mL of floatation supernatant was carefully aspirated to a clean 50mL Falcon tube, resuspended in distilled water and centrifuged again under the same conditions. After discarding the supernatant, the pellet with concentrated eggs was carefully transferred to a 1.5 mL Eppendorf tube, where they were stored at -20°C until DNA isolation.

For centrifugal flotation, ∼2 g of faeces was combined with 10 mL of saturated salt (NaCl, specific gravity = 1.21). The sample was homogenised, passed through a sieve, and centrifuged (800×g for 3 min). Following centrifugation, four drops were collected from the meniscus using the back end of a pipette tip, with two drops transferred to each of two glass slides prepared from the same flotation supernatant. One slide was fitted with a standard coverslip for manual microscopy and the paired slide with an Vetscan Imagyst-compatible coverslip (Zoetis Diagnostics, Australia). The slide with standard coverslip was read on Olympus BX43 Upright Microscope at 40-400× magnification, and subjectively semi-quantitatively recorded presence of parasite eggs from low (+) to high (++++) egg counts. The paired slide was scanned using a Grundium Ocus 40 whole-slide scanner and analysed through the Vetscan Imagyst an AI-enabled image analysis online platform (Zoetis Diagnostics, Australia). Presence with quantity of parasite eggs, cysts and oocysts across the entire slide was reported via online portal. Receiver operating characteristic (ROC) analysis was performed to assess the ability of Vetscan Imagyst egg counts to discriminate samples above predefined FEC thresholds (100, 150, 250, 500 and 1,000 eggs per gram). AUC values and 95% confidence intervals were estimated using the DeLong method. Agreement between diagnostic methods was assessed using Cohen’s kappa coefficient (κ) with 95% confidence intervals and Landis and Koch’s interpretation (0.61-0.80, substantial).

### 2.3. DNA isolation

Genomic DNA extraction was performed using the Isolate II Fecal DNA Kit (Meridian Bioscience, Australia) following the manufacturer’s protocol. Briefly, 750 µL of lysis buffer was added to the pellet tube and disrupted using a FastPrep-24 5G homogeniser (MP Biomedicals, Australia) set to 6.0 m/s for 40 seconds. Blank samples containing only lysis buffer were included following every ten samples to monitor contamination. All samples, including extraction blanks, were eluted in 100 µL of 10 mM Tris-Cl (pH = 8.5) and stored at - 20 °C until further use.

### 2.4. Amplicon deep sequencing and sequence analysis

All DNA samples were subjected to ITS-2 and two isotype-1 β-tubulin fragment amplifications using primer sets modified for Illumina metabarcoding using Next Generation Sequencing (NGS) (Abdullah et al., 2025; Fagundes-Moreira et al., 2026). The ITS-2 target was amplified (NC-1/NC-2) for 25 to 32 cycles, whilst the β-tubulin assays (BZ167: UnAnBZ167-F/R, BZ200: UnAnBZ200-F/R) used 30 to 36 cycles. Each 25 µL PCR mix contained 2 µL of undiluted template DNA (neat) and 400 nM of primers, prepared on a Myra robotic liquid handler with SensiFAST™ SYBR^®^ No-ROX master mix (Meridian Bioscience, Australia). Amplification was performed on a Bio-Rad CFX Opus Real-Time system with an initial denaturation at 95°C (3 min), followed by cycling at 95°C, 57°C and 72°C for 5s, 15s, and 15s, respectively, and a final melt curve analysis. No-template controls (ddH_2_O) were included in every run to detect contamination. Pooled amplicons per individual DNA were indexed, purified and sequenced at the Ramaciotti Centre for Genomics (UNSW, Sydney, Australia) on a MiSeq i100 Plus platform (2×300 bp).

All FastQ files were analysed using the ‘dada2’ package (v1.34) (Callahan et al., 2016) implemented in R v4.4.2 (R Core Team, 2022) to generate amplicon sequence variants (ASVs) and corresponding abundance tables using previously developed pipeline (Abdullah et al., 2025). Primer sequences were removed using ‘cutadapt’ v4.0 (Martin, 2011). Filtering and trimming were performed using filterAndTrim in ‘dada2’ with parameters maxEE = (2,2), truncQ = 2, and maxN = 0. Reads were truncated to 190 bp (forward) and 180 bp (reverse) amplicons, and phiX reads were removed. Followed by ‘dada2’ workflow for filtering, error estimation and denoising. Paired end reads were merged to reconstruct full-length ASVs, and chimera removal as previously described (Abdullah et al., 2025).

Species identification from ITS-2 ASVs was initially performed with IdTaxa algorithm in the ‘DECIPHER’ package (v3.2), against the nemabiome.ca ITS-2 reference database (Workentine et al., 2020). ASVs were subsequently imported into CLC Main Workbench v23 (CLC bio, Qiagen Australia) and top blastn hits against locally curated database was identified. Species identification was confirmed by subsequent phylogenetic reconstruction as each ASV was manually assigned to a species on the tree relative to the reference ITS-2 sequences (KP844736; MT345056; JQ812692; LC036567; JX220891; EU344797; AB793527). For the β-tubulin amplicons (BZ167 and BZ200 assays), species assignment was performed as previously described (Abdullah et al., 2025; Chen et al., 2026; Stocker and Šlapeta, 2025). Briefly, all ASVs were examined in CLC Main Workbench v23 against curated β-tubulin amplicons from *Ancylostoma caninum*, *A. ceylanicum*, *A. duodenale*, *Necator americanus* sourced from ‘WormBase ParaSite’ (Bolt et al., 2018; Harris et al., 2020; Howe et al., 2017) and *Uncinaria stenocephala* (Stocker and Šlapeta, 2025). This comparison enabled identification of exon-intron boundaries and key codons associated with BZ susceptibility or resistance, specifically amino acid positions Q134, F167, E198 and F200 (Venkatesan et al., 2023). Following ASV taxonomical identification and positions coding for amino acid 134, 167, 198 and F200, the ASV were pooled according to parasite species and codons, and raw read counts were converted to relative abundances (percentages). For each sample, the proportional representation of SNPs at codons 134, 167, 198 and 200 was calculated for both β-tubulin amplicons (BZ167, BZ200).

Normality was assessed using the Shapiro-Wilk test. As frequency data were not normally distributed, differences among sampling periods were analysed using the Kruskal-Wallis test. Analyses were performed in GraphPad Prism v11.0.2 (GraphPad Software, Boston, MA, USA).

### 2.5. Data availability

The metadata and intermediate files for all samples are available at LabArchives (https://dx.doi.org/10.25833/ptqs-5j09). Raw FastQ sequence data were deposited at SRA NCBI BioProject: PRJNA1553489.

## 3. Results

### 3.1. High frequency of hookworm-like eggs in shelter samples

Hookworm-like eggs were detected in 70/173 (41%) of canine faecal samples when results from all diagnostic approaches were considered (**Table 1**). Residual canine faecal diagnostic samples (AMH01-AMH175) were received during August 2025 (n=49), December 2025 (n=37), February 2026 (n=45) and March 2026 (n=42). Using faecal egg count, we detected 68 (39%) samples with hookworm-like eggs (average 1,736 eggs per gram, min. 40, max. 24,200). Using Vetscan Imagyst AI-enabled image analysis system following centrifugal floatation, we detected 64 (37%) samples with hookworm-like eggs (average 482 eggs on a slide, min. 1, max. 10,954) and ROC analysis demonstrated good agreement with faecal egg counts for detection of hookworm infections, particularly at higher egg-count thresholds (**Supplementary Data S1**). Using operator read centrifugal faecal floatation, we detected 59 (34%) samples with hookworm-like eggs; subjectively judged as low to high positivity (+: 38, ++: 15, +++: 5, ++++: 1). Fifteen samples had discordant results (**Supplementary Data S2**). The faecal egg count missed two samples that were positive only in one other test: AMH11, Vetscan Imagyst detected 2 eggs; AMH82, flotation detected low (+) hookworm-like eggs. Vetscan Imagyst missed six samples and operator read centrifugal faecal floating missed 11 hookworm-like eggs positive samples. The discordant results had an average faecal egg count of 109 (min. 40, max. 320) and Vetscan Imagyst detected 1 to 7 eggs. Agreement between all faecal egg counts and Vetscan Imagyst was substantial (κ = 0.76, 95% CI 0.56-0.94) as was the agreement between faecal egg count and operator read flotation (κ = 0.61, 95% CI 0.41-0.81).Concentrated hookworm-like eggs from faecal egg count positive samples (n=68) underwent molecular characterisation using ITS-2 rDNA and two β-tubulin fragments (BZ167, BZ200). In addition, 14 samples were reisolated and processed as duplicates (AMH28, AMH34, AMH64, AMH97, AMH134, AMH135, AMH136, AMH137, AMH138, AMH141, AMH143, AMH158, AMH160, AMH161).

**Table 1.** Coproanalysis of hookworm-like eggs from shelter dogs.

|  | Samples (%) | Presence of hookworm-like eggs |  |  |
|---|---|---|---|---|
|  |  | FEC (%) | Vetscan Imagyst (%) | Flotation (%) |
| August | 49 (28) | 19 (39) | 16 (33) | 16 (33) |
| December | 37 (21) | 8 (22) | 8 (8) | 6 (16) |
| February | 45 (26) | 18 (40) | 18 (40) | 17 (38) |
| March | 42 (24) | 23 (55) | 22 (52) | 20 (48) |
| Shelter 1 | 35 (20) | 10 (29) | 8 (23) | 8 (23) |
| Shelter 2 | 96 (55) | 35 (36) | 33 (34) | 31 (32) |
| Shelter 3 | 38 (22) | 20 (53) | 20 (53) | 17 (45) |
| All | 173 | 68 (39) | 64 (37) | 59 (34) |

Note: FEC - faecal egg count.

### 3.2. ITS-2 deep amplicon sequencing reveals predominant *A. caninum* species composition in samples from shelter dogs

Sequencing of the ITS-2 rRNA gene yielded sequencing reads for all 68 samples, but after filtering and taxonomical identification 65 were retained. No hookworm sequence reads were detected in sample AMH04 and two samples (AMH16, AMH24) had <1,000 hookworm sequence reads (206, 13). Based on coproanalysis, AMH04 and AMH24 were previously flagged as discordant (**Supplementary Data S2**). Sample AMH16 and AMH112 included ITS-2 of *Nippostrongylus brasiliensis* that formed 99% and 81% of the sequencing reads, respectively. These sequences were interpreted as environmental contaminants because *N. brasiliensis* is a parasite of rats and is not recognised as a natural infection of dogs. The remaining analysis focused only on hookworms.

Only *A. caninum* and *U. stenocephala* DNA sequences were detected in hookworm positive samples (n=65) with >1,000 good quality hookworm reads; we obtained on average 19,452 good quality hookworm sequence reads (min. 2,748, max. 48,839). All but two samples included *A. caninum* sequence reads, 63/65 (97%). Five samples had *U. stenocephala* sequencing reads (5/65, 8%). Three samples (5%) were coinfected with *A. caninum* and *U. stenocephala* based on ITS-2 reads. Two coinfected samples (AMH12 with FEC = 240, AMH97 with FEC = 120) had 74% and 1% reads of *A. caninum*, respectively, while the remaining belonged to *U. stenocephala*. Coinfected sample AMH112, had only 10 *U. stenocephala* ITS-2 reads (0.36%) belonging to ASV that was however shared with all other *U. stenocephala* positive samples.

### 3.3. Widespread occurrence but low within-sample frequency of the isotype-1 β-tubulin F167Y allele in *A. caninum* hookworm populations from shelter dogs

Sequencing of the isotype-1 β-tubulin gene fragment BZ167 yielded reads from all 68 samples (**Figure 1A**); however, following filtering and taxonomic assignment, 62 samples were retained. No hookworm sequence reads were detected in samples AMH04 and AMH24, both of which had faecal egg counts of 40 eggs per gram (**Table 1**). Four samples (AMH16, AMH22, AMH112 and AMH159) contained fewer than 1,000 hookworm sequence reads (324-892). Based on coproanalysis, four of these six samples had previously been identified as discrepant (**Supplementary Data S2**).

**Figure 1.**
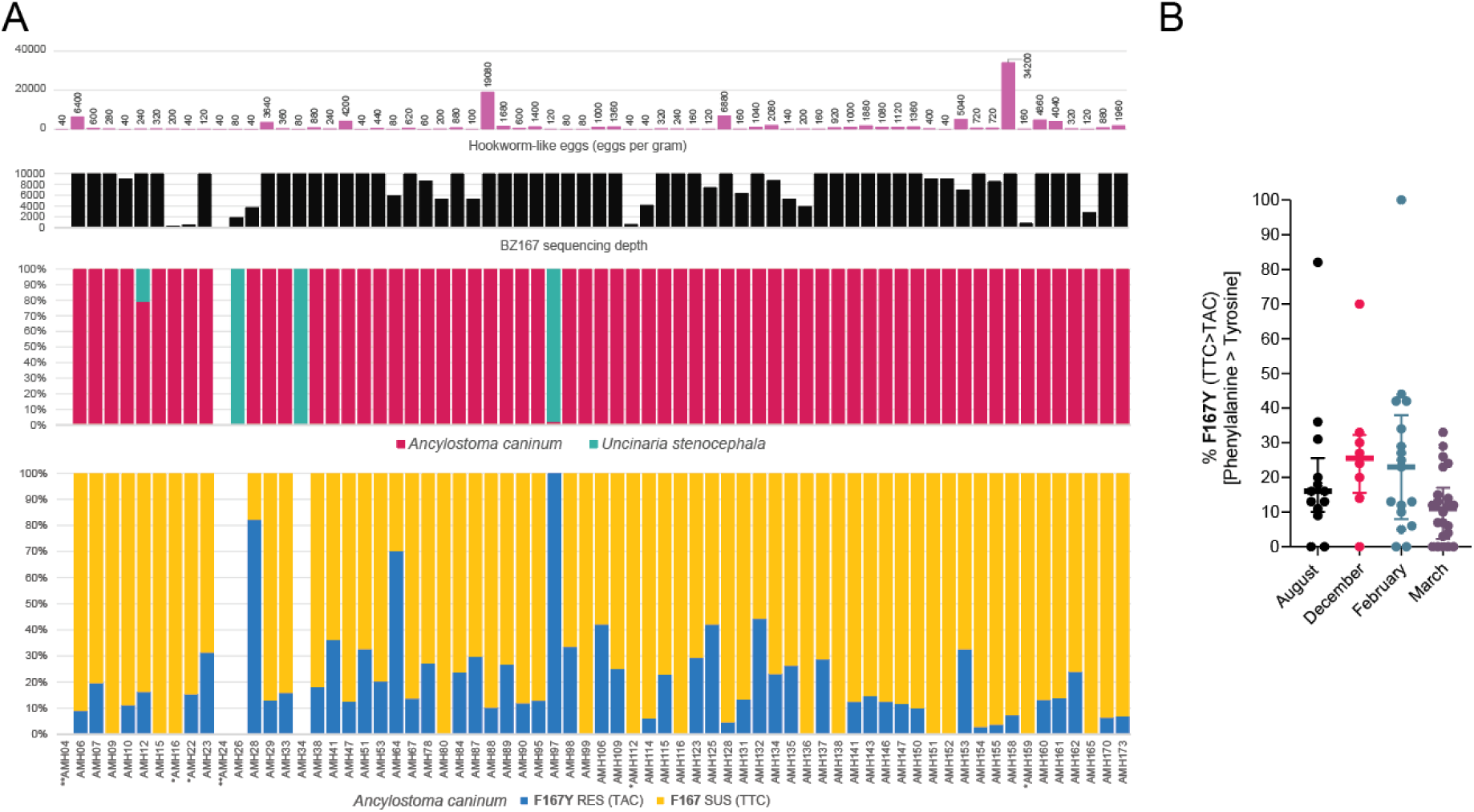
Widespread occurrence of the isotype-1 β-tubulin gene F167Y mutation in shelter-derived hookworm populations. (A) Faecal egg counts (FEC), BZ167 sequencing depth, hookworm species composition and frequencies of the susceptible (F167; TTC) and benzimidazole resistance-associated (F167Y; TAC) alleles in individual samples. (B) F167Y allele frequencies across four sampling periods. Points represent individual *A. caninum*-positive samples and horizontal bars indicate the mean and interquartile range (IQR). F167Y allele frequencies did not differ significantly among sampling periods (Kruskal-Wallis χ² = 7.70, P = 0.053). * Fewer than 1,000 hookworm sequence reads were recovered following filtering and taxonomic assignment. ** No hookworm sequence reads were recovered following BZ167 sequencing.

For the 62 hookworm-positive samples, the BZ167 amplicon yielded a mean of 12,803 high-quality hookworm sequence reads (min. 1,890; max. 34,457). *A. caninum* sequence reads were detected in 60/62 (97%) samples, while *U. stenocephala* sequence reads were detected in 4/62 (7%) samples. Two samples (AMH12, FEC = 240; AMH97, FEC = 120) were coinfected with *A. caninum* and *U. stenocephala*. In these samples, A. caninum represented 79% and 2% of sequence reads, respectively, with the remainder assigned to *U. stenocephala*.

The *A. caninum* isotype-1 β-tubulin gene F167Y mutation was detected in 50/60 (83%) *A. caninum*-positive samples. However, only three samples (6%) contained the mutation at frequencies exceeding 70%. The *A. caninum* isotype-1 β-tubulin gene F167Y mutation was detected with a mean read depth of 2,631 (min. 257, max. 6,567) and a mean allele frequency of 23% (min. 3%, max. 100%) (**Figure 1A**). The mean frequency across the four sampling periods was 20% (95% CI 8-33%), 27% (95% CI 10-44%), 25% (95% CI 12-37%) and 11% (95% CI 7-16%), respectively, and did not differ significantly among groups (Kruskal-Wallis χ² = 7.70, n = 60, P = .053) (**Figure 1B**).

Three samples (AMH28, AMH64 and AMH97) exhibited F167Y allele frequencies exceeding 70%, representing 6% (3/50) of samples carrying the mutation (**Figure 1AB**). Sample AMH97 exhibited 100% F167Y across 267 BZ167 reads; however, this sample was coinfected with *U. stenocephala* and *A. caninum* represented only 2% of hookworm sequence reads. The remaining two samples with >70% F167Y allele frequency (AMH28 and AMH64) were both low egg shedders, with faecal egg counts of 40 and 80 eggs per gram, respectively. Repeated AMH28 and AMH64 samples yielded 45% and 39% F167Y frequency, respectively (**Supplementary Data S3**). Excluding these three samples did not alter the overall pattern, and F167Y allele frequencies remained similar among sampling periods (Kruskal-Wallis χ² = 7.9, P = .116).

The *A. caninum* isotype-1 β-tubulin gene Q134H mutation was rare and was detected in only a single sample (AMH28) at an allele frequency of 7%; repeated sample yielded 11% Q134H frequency.

The *U. stenocephala* isotype-1 β-tubulin gene F167Y mutation was not detected in any of the four *U. stenocephala*-positive samples. The *U. stenocephala* Q134H mutation was detected in one of four samples (25%, 1/4), with an allele frequency of 24% (2,847/11,868 reads; AMH34, 80 eggs per gram); repeated sample yielded 23% Q134H frequency (**Supplementary Data S3**).

Sequencing of the BZ200 fragment encompassing codons E198 and F200 revealed no evidence of BZ resistance-associated variants across all *U. stenocephala* and nearly all *A. caninum* samples. Although the E198A mutation (GAG>GCG) was noted at a low frequency (0.7%; single ASV: 48/6,599 reads) in one *A. caninum* sample (AMH137), the mutation failed to replicate in the duplicate sequenced sample, pointing to a potential artifact (PCR error or sequencing artifact). Consequently, while E198A is a known BZ-resistance marker in ruminant strongyles, its role or presence in hookworms like *A. caninum* is currently unsubstantiated (Dini et al., 2026; George et al., 2022).

### 3.4. Shelter acquisition of *A. caninum* with the isotype-1 β-tubulin gene F167Y mutation

The longitudinal study across four sampling periods enabled us to determine whether dogs sampled on multiple occasions acquired or resolved hookworm infection. We identified a total of 26 dogs that were sampled at least twice during the study period (n = 59 samples; mean 2.3 samples per dog).

Among ten dogs that became newly infected with *A. caninum* during shelter residence, nine (90%, 9/10) acquired hookworm populations carrying the F167Y mutation. Ten dogs (n = 24 samples) were hookworm-negative at their initial sampling but were subsequently positive for *A. caninum* (60-2,080 eggs per gram; **Figure 2**). Nine dogs subsequently shed *A. caninum* carrying the isotype-1 β-tubulin gene F167Y mutation, with within-sample allele frequencies ranging from 4% to 44% (**Figure 2A**).

**Figure 2.**
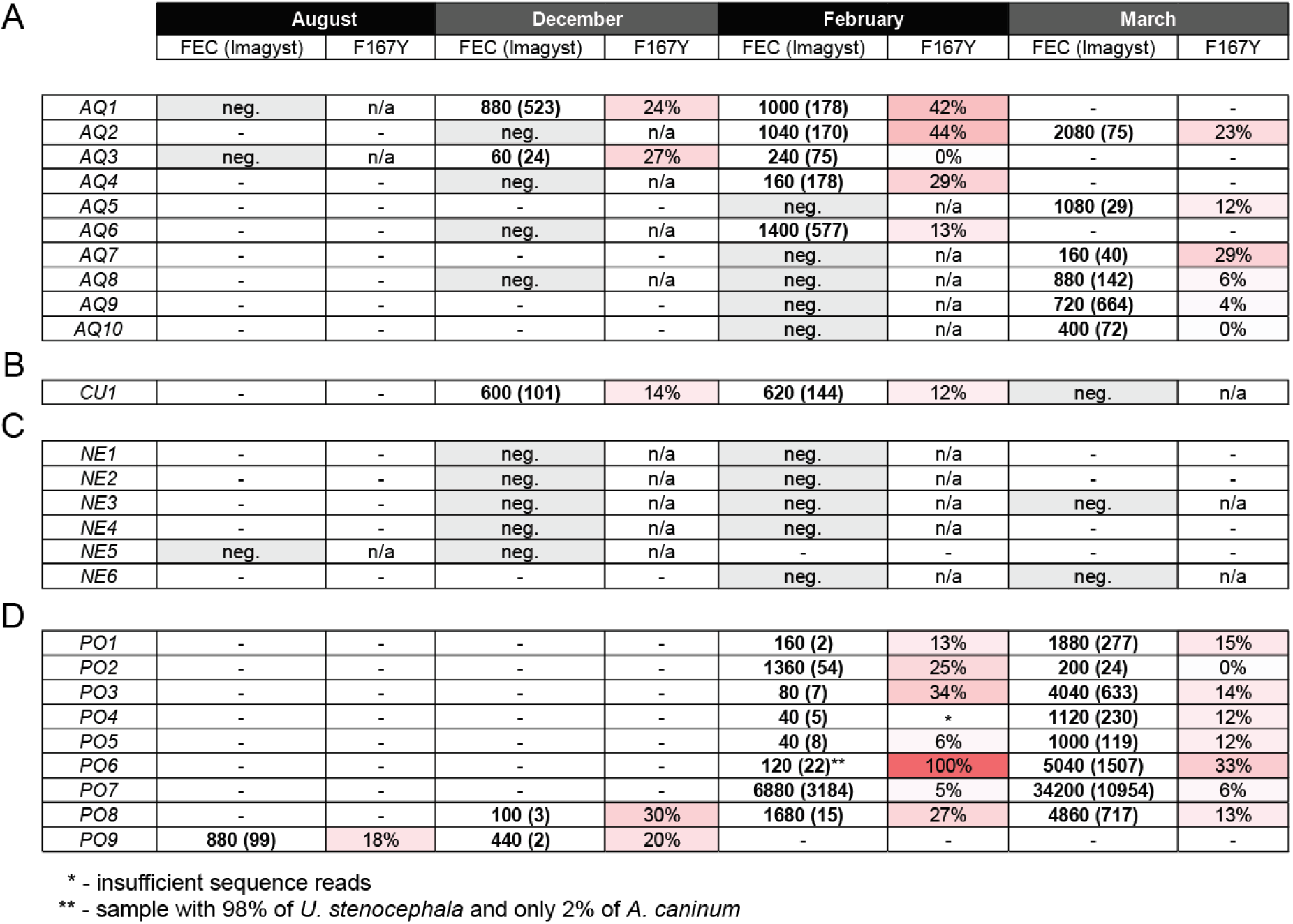
Longitudinal tracking of hookworm infection and the isotype-1 β-tubulin gene F167Y mutation in shelter dogs. Dogs sampled on multiple occasions were grouped according to infection status over time: (A) dogs that acquired *Ancylostoma caninum* infection following an initial hookworm-negative result, (B) dog that resolved hookworm infection during the study period, (C) dogs that remained hookworm-negative throughout the study and (D) dogs that remained hookworm-positive throughout the study. Values represent faecal egg counts (FEC; eggs per gram) with corresponding Vetscan Imagyst egg counts shown in parentheses. F167Y values indicate the within-sample frequency of the *A. caninum* isotype-1 β-tubulin gene F167Y mutation determined by deep amplicon sequencing. Red shading denotes increasing F167Y allele frequency. Negative (neg.), no hookworm eggs detected; n/a, sequencing not performed due to absence of hookworm eggs.

Three dogs remained hookworm-positive over two consecutive sampling occasions, enabling assessment of changes in F167Y allele frequency over time. Dog #AQ1 was tested twice after an initial negative result; the first positive sample (AMH84) had an F167Y allele frequency of 24%, which increased to 42% in the subsequent sample (AMH106). Dog #AQ2 had an F167Y allele frequency of 44% in the first positive sample, followed by 23% in the subsequent sample (AMH132 and AMH134). Dog #AQ3 had an F167Y allele frequency of 27% in the first positive sample, whereas the later sample contained only the susceptible F167 allele (AMH78 and AMH116).

One dog (#CU1; n = 3 samples; **Figure 2B**) was initially positive for hookworms, with F167Y allele frequencies of 14% and 12%, but became hookworm-negative during the study period. Six dogs (n = 13 samples; **Figure 2C**) remained hookworm-negative throughout the study. Nine dogs remained hookworm-positive (n = 19 samples; **Figure 2D**). In one dog, the *A. caninum* F167Y allele frequency declined to 0%, whereas the remaining dogs harboured hookworm populations with F167Y allele frequencies ranging from 6% to 33%.

Dog #PO6 was initially coinfected with *U. stenocephala* (98%) and *A. caninum* (2%) (AMH97), with the *A. caninum* population consisting entirely of the F167Y allele (100%). At the subsequent sampling (AMH153), only *A. caninum* was detected and the F167Y allele frequency was 33%.

## 4. Discussion

The most important finding of this study was that nine of ten dogs that became infected with *A. caninum* during shelter residence subsequently shed hookworms carrying the isotype-1 β-tubulin gene F167Y mutation. This finding supports our hypothesis that dog shelters act as hotspots for the maintenance and dissemination of BZ resistance-associated alleles. The widespread detection of the F167Y mutation across shelter-associated *A. caninum* populations contrasted with its generally low within-sample frequency. A survey of veterinary submissions from the United States reported a lower proportion of samples carrying the F167Y mutation (50%), but a substantially higher proportion of infections dominated by the resistant allele (>70% F167Y frequency; 39% of samples) (Venkatesan et al., 2023). Similarly, a study of veterinary submissions from eastern Australia detected the F167Y mutation in 65% of samples, with 15% exhibiting >70% F167Y frequency (Abdullah et al., 2025). Together, these observations suggest that resistance-associated alleles are broadly disseminated throughout shelter hookworm populations but rarely predominate within individual infections.

Data on persistence of resistant alleles in *A. caninum* under field conditions are currently lacking. Recently, McKean et al. (2026) showed in experimentally infected dogs that serial passaging in the absence of treatment resulted in an unexpected increase in isotype-1 β-tubulin F167Y allele frequency that was associated with increased *in vitro* ivermectin (ML) resistance, but not BZ resistance. Our longitudinal field data did not provide evidence that *A. caninum* isotype-1 β-tubulin F167Y allele frequencies changed consistently over time, nor did we demonstrate ongoing selection within individual dogs. Rather, allele frequencies fluctuated between sampling occasions. Such variation is expected because dogs housed in shelters are continuously exposed to infective larvae, resulting in asynchronous infections composed of parasites acquired at different times.

We hypothesise that the observed within-host allele frequency is determined by two interacting processes. First, continual exposure to infective larvae leads to repeated establishment of new infections, with individual worms differing in developmental stage, time since establishment and onset of patency. Second, allele frequencies are estimated from parasite eggs rather than adult worms. Although deep amplicon sequencing allows analysis of large numbers of eggs, the sampled egg population is not necessarily proportional to the number of adult female worms contributing to it (Šlapeta et al., 2026). Individual female worms differ in fecundity and egg output changes throughout the patency period (McKean et al., 2026). Consequently, some adult worms may contribute disproportionately to the pool of eggs recovered for sequencing. The relative representation of resistant and susceptible alleles within a faecal sample therefore reflects both the composition of the underlying worm population and temporal variation in egg production by individual females. Consequently, the F167Y frequency measured within a single faecal sample is unlikely to represent a stable characteristic of the host and instead reflects the dynamic composition of the underlying parasite population. We therefore consider the average allele frequency observed across the shelter population to provide a more informative estimate of resistant allele persistence within this environmental setting.

It has been proposed that even relatively low BZ resistance allele frequencies within parasite populations may be associated with reduced drug efficacy before overt treatment failure becomes apparent (Kaplan et al., 2026). Our findings demonstrate that the *A. caninum* isotype-1 β-tubulin F167Y mutation is widely embedded within shelter-associated hookworm populations, providing a reservoir from which further selection could occur under sustained BZ exposure. However, the relationship between resistance-associated SNP frequency and clinical treatment outcome remains incompletely understood and the currently recognised β-tubulin mutations should therefore be regarded primarily as markers of resistance risk rather than definitive predictors of treatment success or failure. These observations support ongoing surveillance while highlighting the need for prudent anthelmintic use. Routine mass treatment strategies are unlikely to be sustainable because they may impose strong selection pressure and facilitate accumulation of resistant parasite populations within the environment (Jimenez Castro et al., 2025). Interpretation of routine coprodiagnostic data requires caution, as absence of detected eggs does not equate to absence of infection. In shelter environments, *A. caninum* is maintained not only within infected dogs but also within the contaminated environment that supports ongoing horizontal transmission through percutaneous and oral acquisition of infective larvae.

## 5. Conclusion

The widespread detection of F167Y resistance-associated mutations supports the hypothesis that urban shelters may function as convergence points for resistant *A. caninum* populations. As dogs are subsequently rehomed throughout metropolitan and regional areas, these environments may contribute to the broader dissemination of resistance-associated alleles. The veterinary significance of these dissemination events remains uncertain, as dissemination of resistance-associated alleles does not necessarily translate into persistent transmission or clinically relevant treatment failure. Given the absence of validated molecular markers for other anthelmintic classes, isotype-1 β-tubulin SNPs currently serve as a proxy indicator for broader anthelmintic resistance risk. Continued surveillance and development of targeted anthelmintic stewardship strategies are likely to become increasingly important for dogs housed in high-density environments and for companion dog populations in metropolitan areas.

## Acknowledgements

We thank the Veterinary Pathology Diagnostic Services (VPDS) at The University of Sydney for sharing the residual diagnostic samples. The study was supported by a grant from the Australian Companian Animal Health Foundation (ACAHF) and in part funded by the Sydney School of Veterinary Science Research & Enquiry Unit of Study 2026 funds. This work was completed in partial fulfilment for the requirements of the Doctor of Veterinary Medicine degree, The University of Sydney (AM, SYW).

## References

Abdullah, S., Stocker, T., Kang, H., Scott, I., Hayward, D., Jaensch, S., Ward, M.P., Jones, M.K., Kotze, A.C., Šlapeta, J., 2025. Widespread occurrence of benzimidazole resistance single nucleotide polymorphisms in the canine hookworm, *Ancylostoma caninum*, in Australia. Int J Parasitol 55, 173–182. 10.1016/j.ijpara.2024.12.001.

Bolt, B.J., Rodgers, F.H., Shafie, M., Kersey, P.J., Berriman, M., Howe, K.L., 2018. Using WormBase ParaSite: An integrated platform for exploring helminth genomic data. Methods Mol Biol 1757, 471–491. 10.1007/978-1-4939-7737-6_15.

Callahan, B.J., McMurdie, P.J., Rosen, M.J., Han, A.W., Johnson, A.J., Holmes, S.P., 2016. DADA2: High-resolution sample inference from Illumina amplicon data. Nat Methods 13, 581–583. 10.1038/nmeth.3869.

Chen, Y.J., Li, V., Suwandy, M., Mitrea, I.B., Hayward, D., Jaensch, S., Francis, E.K., Slapeta, J., 2026. Ambient temperature storage in DESS supports molecular studies of benzimidazole resistance from canine hookworm eggs. Int J Parasitol 56, 104745. 10.1016/j.ijpara.2025.10.005.

Dale, A., Xu, G., Kopp, S.R., Jones, M.K., Kotze, A.C., Abdullah, S., 2024. Pyrantel resistance in canine hookworms in Queensland, Australia. Vet Parasitol Reg Stud Reports 48, 100985. 10.1016/j.vprsr.2024.100985.

Dini, M., Chen, R., Redman, E., Leutenegger, C.M., Tereski, J., Loo, S., Savard, C., Jimenez Castro, P.D., Miyashiro, S.I., Gilleard, J., 2026. The detection of three independent benzimidazole resistance mutations in a single *Ancylostoma caninum* population from a Golden Retriever breeding kennel in Sao Paulo, Brazil. Int J Parasitol Drugs Drug Resist 31, 100643. 10.1016/j.ijpddr.2026.100643.

Fagundes-Moreira, R., Bezerra-Santos, M.A., Šlapeta, J., Otranto, D., 2026. Clinical pathology, parasitological evaluation, and anthelmintic resistance assessment in dogs naturally infected with Uncinaria stenocephala. Parasites & Vectors. 10.1186/s13071-026-07688-4.

Geary, T.G., Drake, J., Gilleard, J.S., Chelladurai, J., Castro, P.D.J., Kaplan, R.M., Marsh, A.E., Reinemeyer, C.R., Verocai, G.G., 2025. Multiple anthelmintic drug resistance in the canine hookworm *Ancylostoma caninum*: AAVP position paper and research needs. Vet Parasitol 338, 110536. 10.1016/j.vetpar.2025.110536.

George, S., Suwondo, P., Akorli, J., Otchere, J., Harrison, L.M., Bilguvar, K., Knight, J.R., Humphries, D., Wilson, M.D., Caccone, A., Cappello, M., 2022. Application of multiplex amplicon deep-sequencing (MAD-seq) to screen for putative drug resistance markers in the *Necator americanus* isotype-1 beta-tubulin gene. Sci Rep 12, 11459. 10.1038/s41598-022-15718-1.

Harris, T.W., Arnaboldi, V., Cain, S., Chan, J., Chen, W.J., Cho, J., Davis, P., Gao, S., Grove, C.A., Kishore, R., Lee, R.Y.N., Muller, H.M., Nakamura, C., Nuin, P., Paulini, M., Raciti, D., Rodgers, F.H., Russell, M., Schindelman, G., Auken, K.V., Wang, Q., Williams, G., Wright, A.J., Yook, K., Howe, K.L., Schedl, T., Stein, L., Sternberg, P.W., 2020. WormBase: a modern model organism information resource. Nucleic Acids Res 48, D762–D767. 10.1093/nar/gkz920.

Howe, K.L., Bolt, B.J., Shafie, M., Kersey, P., Berriman, M., 2017. WormBase ParaSite - a comprehensive resource for helminth genomics. Mol Biochem Parasitol 215, 2–10. 10.1016/j.molbiopara.2016.11.005.

Jimenez Castro, P.D., Howell, S.B., Schaefer, J.J., Avramenko, R.W., Gilleard, J.S., Kaplan, R.M., 2019. Multiple drug resistance in the canine hookworm *Ancylostoma caninum*: an emerging threat? Parasit Vectors 12, 576. 10.1186/s13071-019-3828-6.

Jimenez Castro, P.D., Willcox, J.L., Rochani, H., Richmond, H.L., Martinez, H.E., Lozoya, C.E., Savard, C., Leutenegger, C.M., 2025. Investigation of risk factors associated with *Ancylostoma* spp. infection and the benzimidazole F167Y resistance marker polymorphism in dogs from the United States. Int J Parasitol Drugs Drug Resist 27, 100584. 10.1016/j.ijpddr.2025.100584.

Kaplan, R.M., Jimenez Castro, P.D., Marsh, A.E., Jesudoss Chelladurai, J.R.J., Gilleard, J.S., 2026. American Association of Veterinary Parasitologists Hookworm Task Force review on best practices for diagnosis and treatment of multidrug-resistant hookworms. J Am Vet Med Assoc 264, 1214–1222. 10.2460/javma.26.02.0094.

Kitchen, S., Ratnappan, R., Han, S., Leasure, C., Grill, E., Iqbal, Z., Granger, O., O’Halloran, D.M., Hawdon, J.M., 2019. Isolation and characterization of a naturally occurring multidrug-resistant strain of the canine hookworm, *Ancylostoma caninum*. Int J Parasitol 49, 397–406. 10.1016/j.ijpara.2018.12.004.

Leutenegger, C.M., Evason, M.D., Willcox, J.L., Rochani, H., Richmond, H.L., Meeks, C., Lozoya, C.E., Tereski, J., Loo, S., Mitchell, K., Andrews, J., Savard, C., 2024. Benzimidazole F167Y polymorphism in the canine hookworm, *Ancylostoma caninum*: Widespread geographic, seasonal, age, and breed distribution in United States and Canada dogs. Int J Parasitol Drugs Drug Resist 24, 100520. 10.1016/j.ijpddr.2024.100520.

Martin, M., 2011. Cutadapt removes adapter sequences from high-throughput sequencing reads. Embnet.journal, 17, 10–12. 10.14806/ej.17.1.200.

McKean, E.L., Jackson, C.A., O’Halloran, D.M., Hawdon, J.M., 2026. Association between macrocyclic lactone and benzimidazole resistance in the dog hookworm *Ancylostoma caninum*. J Parasitol 112, 389–396. 10.1645/25-80.

R Core Team 2022. The R Project for Statistical Computing (Vienna, Austria, R Foundation for Statistical Computing).

Raza, A., Rand, J., Qamar, A.G., Jabbar, A., Kopp, S., 2018. Gastrointestinal parasites in shelter dogs: Occurrence, pathology, treatment and risk to shelter workers. Animals (Basel) 8. 10.3390/ani8070108.

Šlapeta, J., Krucken, J., Rojas, A., Chambers, A., Melville, L.A., Martinez-Valladares, M., Canton, C., Francis, E.K., Zahid, O., Albuquerque, A.C.A., Bartley, D.J., Bassetto, C.C., Byrne, O., Colella, V., Costa-Junior, L.M., Doyle, S.R., Evans, M., Ghafar, A., Godoy, P., Hayashi, N., Helal, M.A., Huggins, L.G., Jabbar, A., Jones, R.A., Karani, B.E., Liron, J.P., Mate, L., McEvoy, A., Mohammedsalih, K.M., Mulcahy, G., Nielsen, M.K., Pafco, B., Peachey, L.E., Robleto-Quesada, J., von Samson-Himmelstjerna, G., de Sousa-Paula, L.C., Gilleard, J.S., 2026. Ten simple rules for implementing deep amplicon sequencing in parasitology. Int J Parasitol 56, 104751. 10.1016/j.ijpara.2025.11.003.

Stocker, T., Abdullah, S., Scott, I., Šlapeta, J., 2026. Genome skimming of dog faecal samples reveals mitogenomes indistinguishable from those of red fox-derived *Uncinaria stenocephala*. Parasitology 153, 1–8. 10.1017/S0031182026101565.

Stocker, T., Šlapeta, J., 2025. Characterisation of beta-tubulin isotypes in *Uncinaria stenocephala* and implications for benzimidazole resistance in hookworms. Vet Parasitol 339, 110569. 10.1016/j.vetpar.2025.110569.

Venkatesan, A., Jimenez Castro, P.D., Morosetti, A., Horvath, H., Chen, R., Redman, E., Dunn, K., Collins, J.B., Fraser, J.S., Andersen, E.C., Kaplan, R.M., Gilleard, J.S., 2023. Molecular evidence of widespread benzimidazole drug resistance in *Ancylostoma caninum* from domestic dogs throughout the USA and discovery of a novel beta-tubulin benzimidazole resistance mutation. PLoS Pathog 19, e1011146. 10.1371/journal.ppat.1011146.

Workentine, M.L., Chen, R., Zhu, S., Gavriliuc, S., Shaw, N., Rijke, J., Redman, E.M., Avramenko, R.W., Wit, J., Poissant, J., Gilleard, J.S., 2020. A database for ITS2 sequences from nematodes. BMC Genet 21, 74. 10.1186/s12863-020-00880-0.

