## Supplementary Data S3 for "Dog shelter facilitated transmission and maintenance of benzimidazole resistance-associated alleles in canine hookworms"

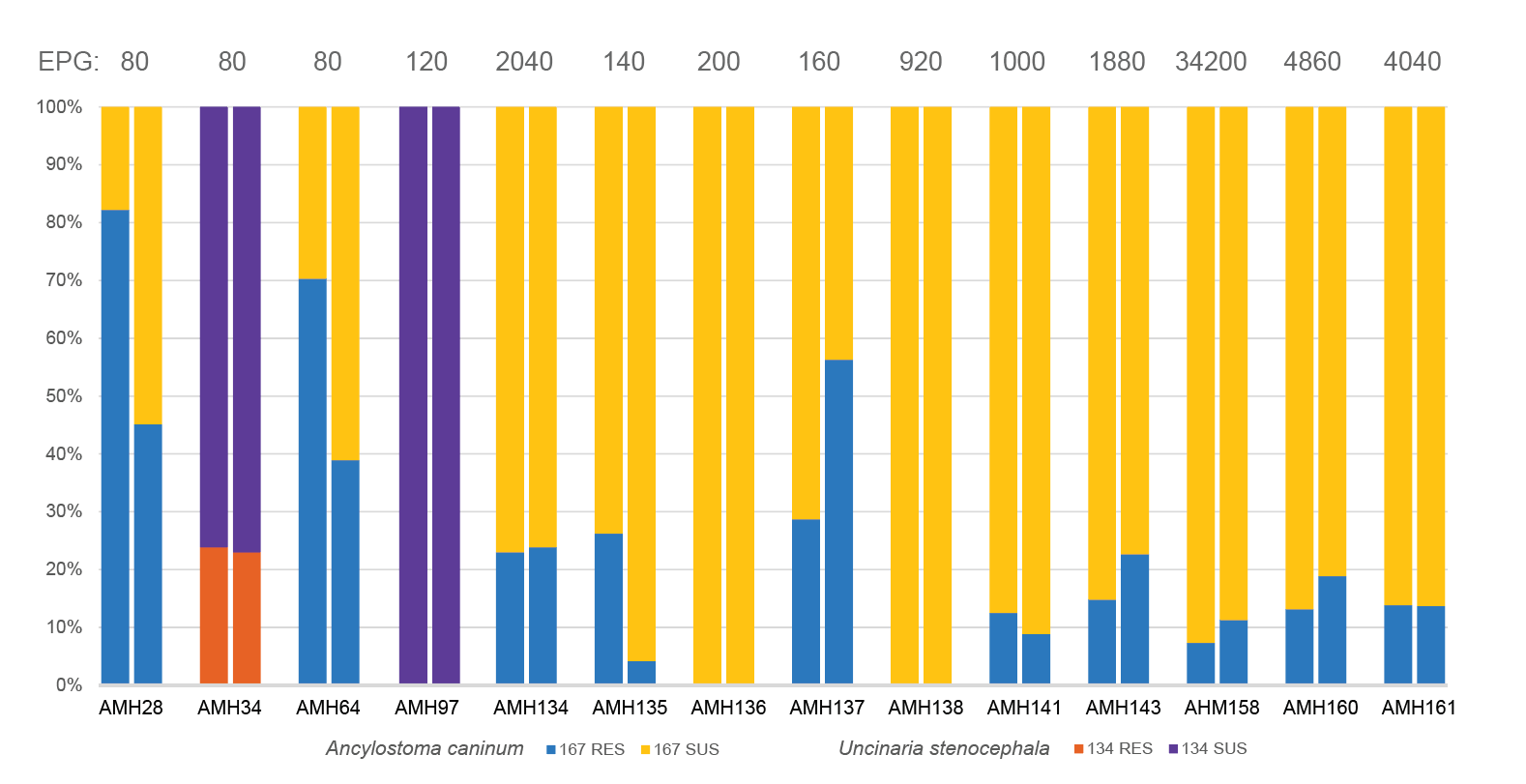

**Supplementary Figure S1.** Replicate BZ167 deep amplicon sequencing of hookworm-positive samples. Duplicate aliquots of hookworm eggs from the same faecal sample were independently reisolated, amplified and sequenced using the BZ167 isotype-1 β-tubulin assay. Bars show frequencies of susceptible and resistance-associated alleles at codon 167 in *Ancylostoma caninum* (F167Y) and codon 134 in *Uncinaria stenocephala* (Q134H). Numbers above bars indicate faecal egg counts (FEC; eggs per gram). Replicate samples are presented side-by-side to illustrate variation in allele-frequency estimates among independently processed egg aliquots from the same faecal sample. Samples AMH34 and AMH97 contained *U. stenocephala* and are therefore represented by codon 134 allele frequencies.
